# Innate attraction and aversion to odors in locusts

**DOI:** 10.1101/2023.04.05.535770

**Authors:** Subhasis Ray, Kui Sun, Mark Stopfer

## Abstract

Many animals display innate preferences for some odors, but the physiological mechanisms underlying these preferences are poorly understood. Here, with behavioral tests, we establish a model system well suited to investigating olfactory mechanisms, the locust *Schistocerca americana*. We conducted open field two-choice tests with purely olfactory stimuli. In these tests, newly hatched locusts navigated toward, and spent time near, the source of a food odor blend, crushed wheat grass. In similar tests, we found that hatchlings avoided moderate concentrations of major individual components of the food blend odor, 1-hexanol and hexanal. They were neither attracted nor repelled by a lower concentration of 1-hexanol, but were moderately attracted to a low concentration of hexanal. These results establish that hatchlings have a strong, innate preference for food odor blend, but the valence of the blend’s individual components may be different and may change depending on the concentration. This suggests innate odor preferences may emerge from more complex processing pathways than labeled lines. Our results provide a useful entry point for an analysis of physiological mechanisms underlying innate sensory preferences.

## Introduction

Animals are born with some innate sensory and behavioral preferences which develop further with age and experience. In humans, newborn females are attracted to odors they are exposed to immediately after birth (Balogh and Porter, 1986), and neonates of both sexes can learn to associate an artificial odor with their mother in as little as one week (Schleidt and Genzel, 1990). Rabbit pups, in contrast to humans, receive little maternal care, so their survival after birth depends on their ability to successfully locate mother’s nipple to obtain milk. It has been shown that newborn rabbits use pheromonal cues for nipple-searching behavior without requiring any postnatal learning (Hudson, 1985; Schaal et al., 2003). Insects such as locusts, whose eggs are laid in sand, do not usually hatch in contact with food; nor do they receive any parental care. So, one might hypothesize these animals must come equipped with innate sensory capacities enabling them to navigate to food, perhaps, at least in part, by tracking food odors. Most food odors are conveyed by combinations of volatile chemicals emanating from the food. For example, wheat grass, which is eagerly eaten by locusts, releases at least 18 different volatiles, with 1-hexanol as the dominant component (Eissa et al., 2018).

Here we investigated the intrinsic preference of newly hatched locusts for a complex food odor by using a low-cost open field setup to conduct a two-choice behavior test. Monitoring behavior in locusts in open field settings can be challenging because even newly hatched locusts can jump and make unpredictable changes in direction. Moreover, their movements can be very intermittent, requiring lengthy experiments to register their choices. Therefore, we developed a new software toolkit for recording and tracking multiple animals (Ray and Stopfer, 2022). Using this toolkit, we recorded and analyzed the movements of hundreds of newly hatched locusts under various conditions. We found that even without any prior exposure to food, hatchlings could navigate to the source of a complex food odor blend, wheat grass juice. Notably, while attracted to the complex food blend, hatchlings were repelled by even low concentrations of the major component of the blend, 1-hexanol, presented alone. We also found the hatchlings were attracted by low concentrations of hexanal, which is both a food blend component and an aggregation pheromone but avoided it at higher concentrations. Thus, our results show that naïve hatchlings are innately attracted by food odor, that they can navigate to its source, and that individual monomolecular components of an attractive food odor can change valence depending on concentration.

## Materials and Methods

### Behavior arena

The behavior arena (Figure 1) consisted of a clear acrylic tray (30.5 cm x 30.5 cm x 5 cm) with a non-reflective glass (33 cm x 33 cm, True Vue) cover (Figure 1a). To reduce visual clutter from reflections, the inner walls of the tray were covered with window film. Air was drawn through the arena through two nylon mesh-covered slots on opposite walls by a computer fan (Comair Rotron 25 mm x 10 mm 12V DC) powered by a DC adapter with adjustable voltage output. We used a smoke test to visualize and calibrate the airflow, adjusting the fan’s supply voltage to minimize turbulence, resulting in an air speed of about 0.35 m/s in front of the exhaust slot.

**Figure 1:**
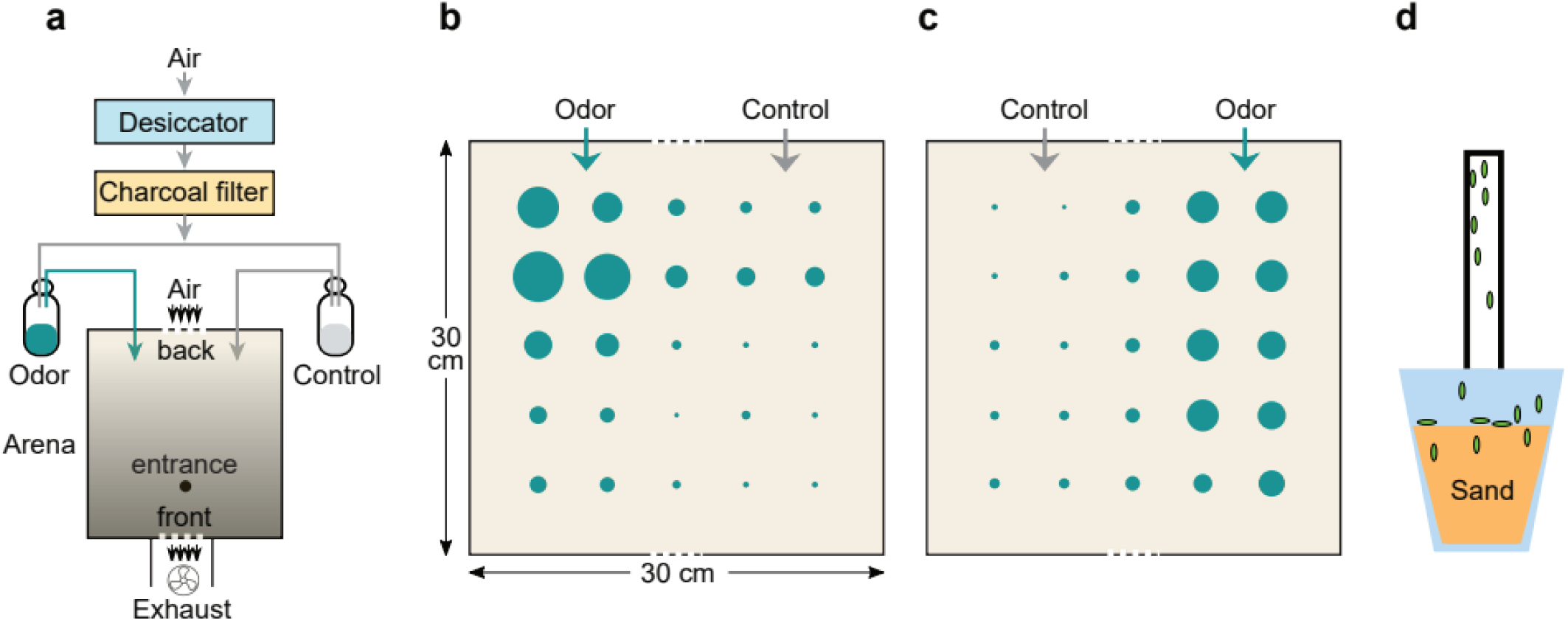
(a) Experimental setup for testing odor preference in insects; (b) and (c) normalized odorant concentration measured by a mini PID at different locations of the arena when delivered through the left and the right port, respectively. (d) Technique for collecting hatchlings without touching them.

The arena was illuminated from below by a tube light placed near the back of the arena. The bottom of the arena was covered with white paper on the outside, which diffused the light and created a gradient so the back of the arena was brighter than the front. After entering near the front of the arena, hatchlings tended to phototax towards the back. A cellphone based light meter confirmed the left and right sides near the back of the arena were equally bright.

### Odor delivery

Two 50 ml glass bottles were fitted with airtight silicone plugs pierced by two polyethylene tubes (Figure 1a). Desiccated, charcoal filtered air flowed into each bottle through one tube and out of the other tube. These outlets were inserted into the behavior arena through holes drilled on the airflow-entry wall, opposite from the exhaust fan. Odor and control ports were alternated between experiments.

### Odorants

To test responses to a food odor, we used wheat grass grown in our laboratory. Fresh grass was crushed with a mortar and pestle, and the juice was then pressed through a cell filter mesh. 1 ml extracted juice was placed as a test odorant in one bottle, and 1 ml deionized water in the other bottle served as control. Bottles were thoroughly cleaned and dried between trials.

To test monomolecular odorants, we used 1-hexanol (Sigma), the dominant component of grass juice volatiles, and hexanal (Sigma), which is also a significant component of grass juice (Eissa et al., 2018). These odorants were diluted in mineral oil, and pure mineral oil was used as the control. All concentrations (see Results) are reported as volume / volume.

### Odor intensity map in arena

To determine the distribution of odorant within the arena, we used a photo-ionization detector (PID, Aurora Scientific) to measure odorant intensity in a 5 cm square grid (Figure 1B). 100% 1-hexanol was delivered through the port on one side as described above, with the glass cover in place. A polyethylene tube attached to the PID was inserted through the entrance (Figure 1a) and placed at a grid point, and the PID signal was recorded thrice for 10 s at 20 s intervals. The baseline signal of the PID was also measured without the odorant. Figures **1b** and **c** show, for odor delivered on the left and the right respectively, the time-averaged odor signal from the second trial, normalized by the time-averaged baseline signal [(odor – baseline) / (max(odor) – baseline)]. These measurements confirmed that odorants were laterally distributed, as desired.

### Behavior experiments

American desert locusts (*Schistocerca americana*) reared in our crowded laboratory colony laid eggs in sand-filled plastic cups. These egg cups were cleaned of plant and other matter and were then kept in an incubator at about 29 °C until hatching. Cups with fresh hatchlings were fit with a plastic test tube through the lid and wrapped with a black cloth (Figure 1d). Locusts tend to climb upwards (negative geotaxis) and towards light (positive phototaxis). The hatchlings therefore spontaneously climbed up into the test tube. When 10-20 hatchlings had climbed up, the test tube was removed and attached to the entrance of the arena from below. The entrance was kept covered with a small petri dish. To start the experiment, the petri dish was removed, allowing hatchlings to enter the arena. This procedure eliminated the need to handle the hatchlings before testing their behavior.

To test whether the antennal olfactory system was needed to navigate within the arena, hatchlings collected in a test tube were anaesthetized by cooling in ice, and then, under a dissection microscope, both antennae were removed with a pair of sharp micro scissors and fully covered with wax to maximize sensory elimination. Hatchlings then recovered at room temperature for about 30 minutes in a test tube before being introduced to the behavior arena as described above.

To test whether cooling the hatchlings affected their behavior, animals were cooled in ice as above, but their antennae were left intact. These hatchlings were also allowed to recover at room temperature for about 30 minutes before they were introduced to the arena.

After each trial in the two-choice behavior tests, the inside of the arena was wiped with 70% ethanol and allowed to dry completely. A new batch of animals was used for each trial.

### Behavior tracking

The behavior of the locusts in the arena was recorded by a USB web camera (Logitech Pro Stream C922x) for 100-minute segments using the Capture utility of the Argos toolkit (Ray and Stopfer, 2022). This utility later processed segments offline, deleting portions in which no movement had been detected. The reduced videos were then processed by the Argos Tracking tool to automatically track hatchling movements. To accurately map tracks in all videos to the same coordinates, the four corners of the arena floor were manually marked in each video, and then based on these points, the transformation matrix from video coordinates to world coordinates was computed. The tracks were then transformed by this matrix from video coordinates into world coordinates. Finally, because the odor and control ports were alternated between trials, they were brought into the same alignment by flipping the coordinates left-right as necessary.

### Data Analysis

All data analysis was carried out by custom scripts written in Python using Python-scipy stack including the Pandas library.

We used t-tests to compare stay times, i.e., the total time a hatchling spent in a Region of Interest (ROI) and ROI entries (total number of times a hatchling crossed into an ROI) between odor and control sides. Although none of the differences were normally distributed (confirmed by Shapiro-Wilk tests), the use of t-tests was appropriate because n > 30 for all comparisons, and thus the central limit theorem could be applied. We also conducted nonparametric Wilcoxon signed rank tests which produced p values (not shown) supporting the same conclusions reported here.

To compare attraction toward the test odor port and the control port over time we used the lifelines module in Python for survival analysis with Kaplan-Meier fit. The Kaplan-Meier curve is a step function indicating the probability of “survival,” i.e. the probability that an event of interest has not yet occurred. In the experiments comparing attraction towards grass juice odor and control, we computed, for each hatchling, the interval between first detecting the hatchling in the arena and its first entrance to the ROI around the odor port. We used these intervals to make a survival plot, with the entry into the ROI as the event of interest (orange line). Animals which did not enter the ROI within the experiment duration of 100 minutes were “censored.” Similarly, the first entry into the ROI around the control port was the event of interest for control, and the time interval from first detecting the hatchling until this event was used in the Kaplan-Meier fit (blue line). The log-rank test was used to compare these curves. Note that here we are comparing the first entry into the odor ROI with that into the control ROI, and they are not mutually exclusive; if a hatchling visited both ROIs, then it is included in both curves with the corresponding time intervals.

## Results

### Naïve locusts are attracted to food odor

To determine whether locusts are innately attracted to food odor, we conducted a two-choice test between grass juice odor and water vapor. We used newly hatched locusts that had no prior exposure to food or food odors (see Materials and Methods). In this first instar stage the hatchlings have not yet developed wings and move by walking or jumping. Figure 2A shows overlaid tracks from all hatchlings in all experiments. Track color in these images was set to change from dark purple to blue to green to yellow over time, showing that many hatchlings moved directly towards the source of grass juice odor after entering the arena. To quantify the affinity of the hatchlings for the food odor, we computed the amount of time they spent in a semicircular region of interest (ROI) of 4 cm radius around each port. Hatchlings spent significantly more time near the odor port than the control port (Figure 2b; paired t-test, n=213, p=6.44e-11). We also found the hatchlings crossed into the food odor port ROI significantly more times than into the control port ROI (Figure 2c, paired t-test, n=213, p=1.16e-6), indicating that they returned to the food odor source again and again while exploring the arena. We computed a preference index (PI) based on total stay time in the ROIs as (∑t_Odor_ – ∑t_Control_) / (∑t_Odor_ + ∑t_Control_), for sums over the entire population, of PI = 0.41. We also computed a PI for the number of ROI entries using the same formula, replacing time *t* with the number of ROI entries *n*, and this yielded PI = 0.32. Both PIs indicate the hatchlings move preferentially toward grass juice odor.

**Figure 2:**
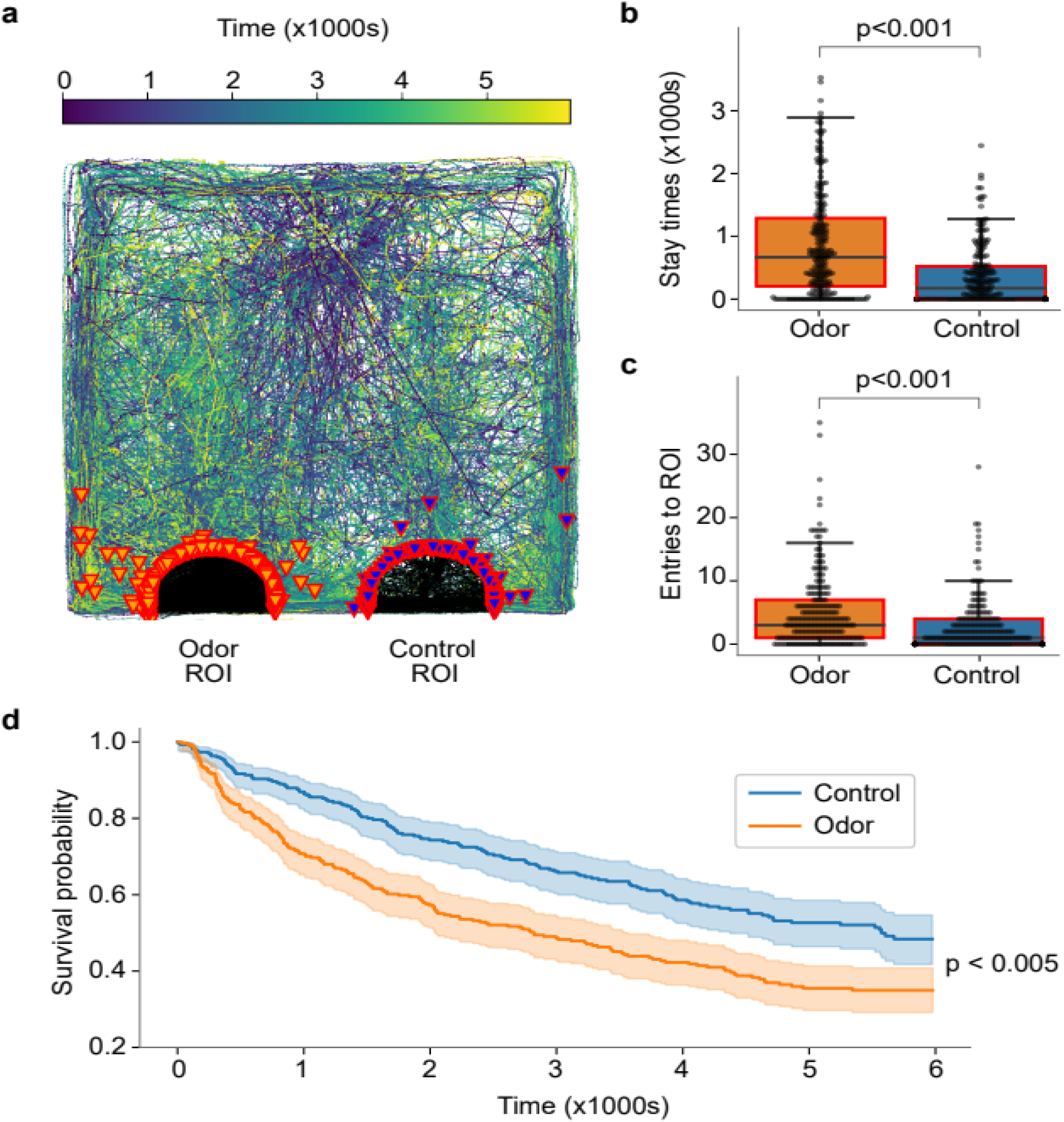
(a) Overlaid tracks of all hatchlings in test of attraction toward wheat grass juice. Purple/dark blue – early part of the track, light green/yellow – late part of the track. Portions of tracks within ROI marked in black, orange arrow heads indicate hatchlings crossing into the ROI around grass juice and magenta arrow heads indicate hatchlings crossing into the ROI around control. (b) Hatchlings stay within the ROI around grass juice (orange) longer than that around control (blue). (c) The number of entries into the odor ROI is also larger than control, indicating that the hatchlings cross into it more often. (d) Kaplan-Meier survival plot, where events are defined as first entries into the ROI around odor (orange) or control (blue), shows hatchlings quickly head toward the odor port.

Finally, we applied a survival analysis using the Kaplan-Meier (KM) method to determine the rates at which hatchlings moved toward odor or control ports (log-rank test statistic=20.17, p < 0.005, see Methods). In this statistic, hatchlings are removed from the “survival” pool when they first enter either ROI, so survival curves shown in Figure 2d indicate the time it took for each hatchling to choose. The KM plots show hatchlings entered the food odor ROI significantly faster than the control ROI. Also, the slope of the curve for the odor is steeper at the beginning, indicating that the hatchlings tend to move quickly towards the odor port upon entering the arena. Because no visual or other cues distinguished the source of the food odor from that of the control, this result establishes that hatchlings could use odor guided navigation to reach the source of an attractive food blend odor.

### Attraction to food odor is mediated by antennae

Antennae are the main olfactory organs in insects, but locusts also have odorant receptors on their mouth parts (Keil, 1999). To test whether the antennal olfactory pathway was necessary for the hatchlings’ innate attraction toward food odor, we conducted behavioral experiments identical to those described above but with hatchlings whose antennae had been removed while they were anesthetized by cooling (see Materials and Methods). We found hatchlings lacking antennae were equally attracted to food odor and control ports (Figure 3a-c; two-tailed paired t-test, for stay time n = 35, p = 0.44, PI = -0.10, and for number of ROI entries n=35, p= 0.57, PI=0.07). We noted that these hatchlings tended to follow circular paths while heading toward the back of the arena, possibly drawn there by the brighter light or by air movements; earlier experiments with other insects where a single antenna was cut did not produce circling behavior (Schultheiss et al., 2020).

**Figure 3:**
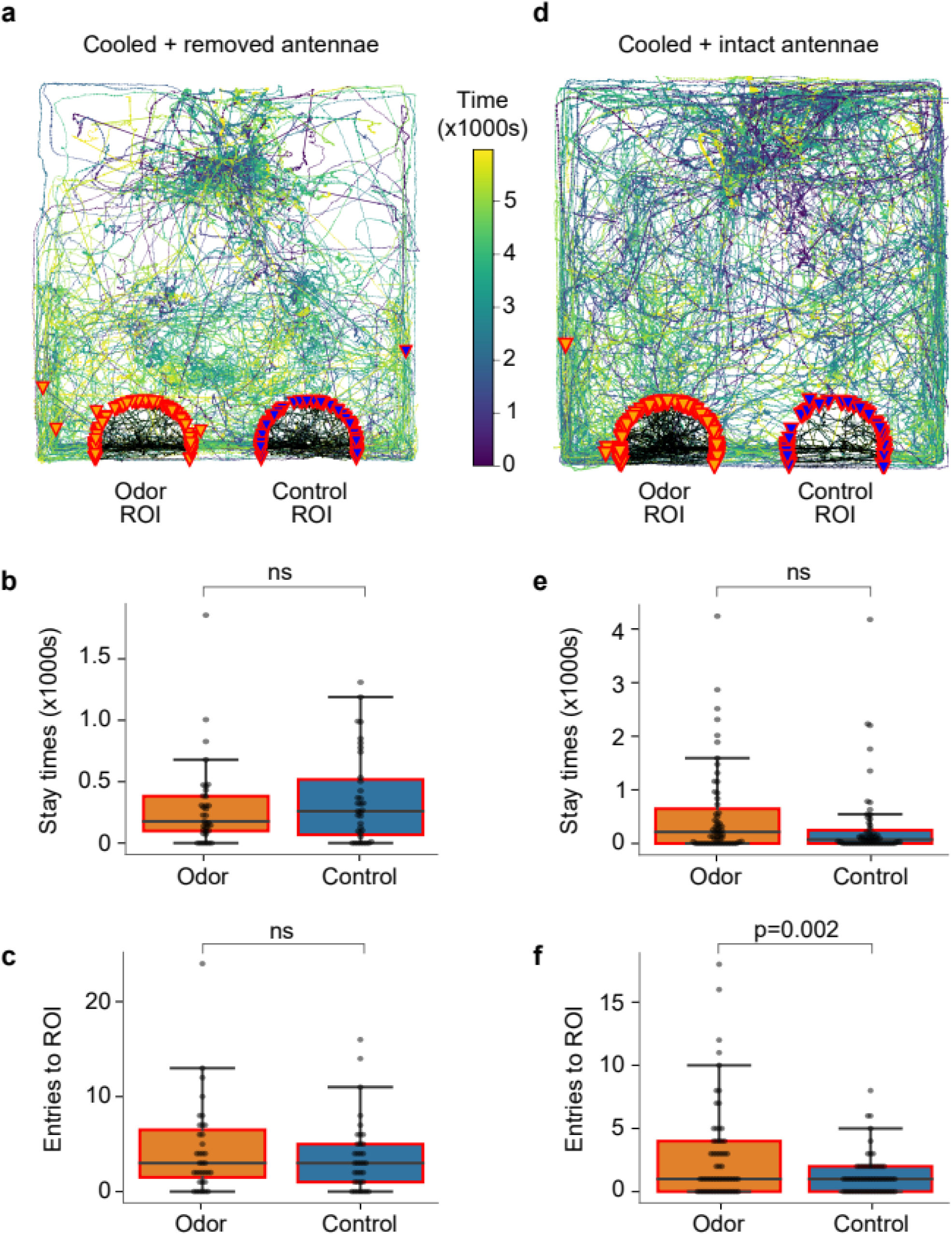
Hatchlings with their antennae removed do not show any difference in attraction towards grass juice odor and control. (a) Overlaid tracks from all hatchlings tested; notably, some, lacking antennae, walked in circular patterns. (b) Stay time in control ROI and odor ROI, (c) number of entries into control ROI and odor ROI. Control experiments revealed that anesthesia by cooling does not in itself affect odor preference and navigation to a preferred odor source when antennae are intact: (d) overlaid tracks of all the hatchlings from these control experiments, (e) stay time in control ROI and odor ROI; results fell just short of statistical significance, (f) number of entries into control ROI and odor ROI.

To test for possible effects of anesthesia by cooling, we also conducted control experiments in which hatchlings were cooled over ice, but their antennae were left intact. These animals, after recovering at room temperature, were, by most measures, significantly attracted towards food odor (Figure 3d-f; paired t-test, for stay-time n = 59, p=0.08, PI = 0.26, and for the number of ROI entries n = 59, p = 0.002, PI = 0.38).

Together, these results indicate that the antennal olfactory pathway primarily mediates the hatchlings’ naïve attraction toward food odor.

### Naïve hatchlings avoid a single component of food odor blend

We next asked whether single components of grass odor attract the hatchlings. The major monomolecular volatile released by wheat grass is 1-hexanol (Eissa et al., 2018). Humans perceive this chemical as smelling like freshly cut grass. Also, electrophysiological experiments have shown this odorant evokes strong neural activity in many projection neurons in the locust antennal lobes (Stopfer et al, 2003). We therefore conducted two-choice tests as above but replaced the food blend odorant with 1-hexanol diluted in mineral oil (1% v/v), tested against pure mineral oil. Notably, the hatchlings were not attracted to 1-hexanol, and instead showed a significant tendency to avoid it (Figure 4a): hatchlings spent significantly less time within the ROI around the odor port than the control port (Figure 4b, paired t-test: n=123, p=3.96e-4, PI= -0.38), and made fewer entries to the odor port than the control port (Figure 4c, paired t-test: n = 123, p=9.05e-5, PI= -0.54).

**Figure 4:**
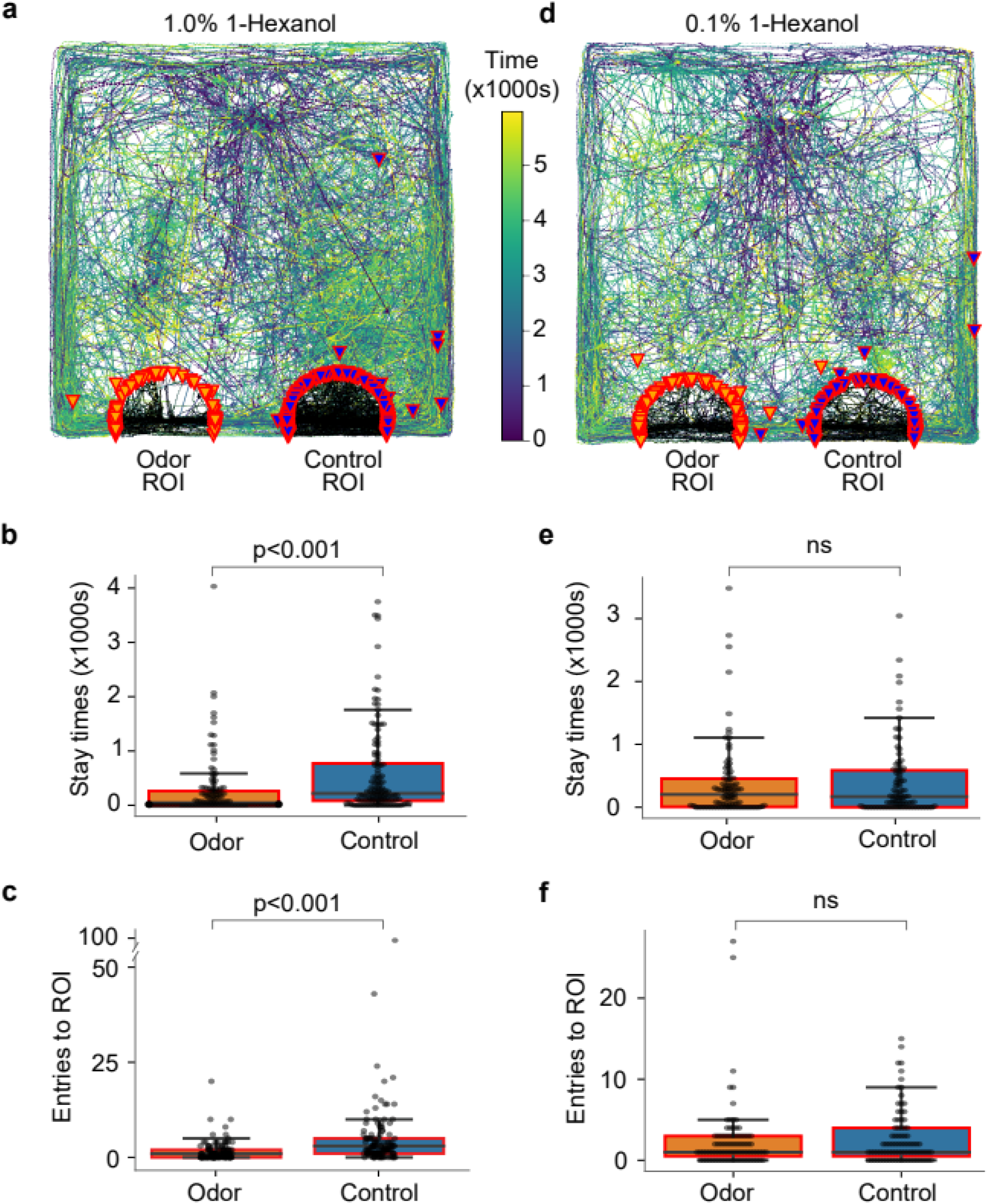
Naive hatchlings avoid 1-hexanol (a) Overlaid tracks of hatchlings tested for affinity towards 1% 1-hexanol compared to mineral oil. (b) Stay-time of hatchlings in the ROI around 1-hexanol compared to that around control. (c) Number of ROI entries. (d) Overlaid tracks of hatchlings tested with 0.1% 1-hexanol. (e) The stay-time in the odor ROI was not significantly different from that around control. (f) Number of ROI entries did not show significant difference between odor and control, either.

Animals are often repelled by strong odors (Laing et al., 1978; Semmelhack and Wang, 2009), so we next reduced the concentration of 1-hexanol in the odor bottle by an order of magnitude (0.1% v/v). However, this concentration of 1-hexanol elicited neither attraction nor repulsion from hatchlings, which showed no difference in affinity for the odorant or the control (Figure 4d-f). In this case the PI for stay time was -0.02 and that for the number of ROI entries was -0.06. Two-tailed paired t-tests showed no differences (n = 87, p = 0.86 for stay time; n = 87, p = 0.60 for number of ROI entries). Together, these results indicate hatchlings were not attracted by 1-hexanol alone.

### Attraction to an aggregate pheromone depends on its concentration

Another major component of wheat grass juice is hexanal (Eissa et al., 2018), which has also been identified as a component of the aggregation pheromone blend in a closely related locust species, *Schistocerca gregaria* (Torto et al., 1996). We found dilute hexanal in the odor bottle (0.9% v/v in mineral oil) repelled hatchlings (Figure 5 a-c; n = 115, paired t-test, p = 2.93e-4, PI = -0.27 for stay-time and p = 1.80e-4, PI = -0.34 for number of ROI entries). Notably, further diluted hexanal (0.225% v/v) attracted hatchlings as suggested by significantly longer stay times near its outlet (Figure 5d-f; n = 95, paired t-test, p = 0.038 and PI = 0.14), though the difference in the number of entries into ROI fell short of statistical significance despite showing a positive preference index (p = 0.15, PI = 0.21). Together, these results demonstrate that even an ethologically important attractive odor can have the opposite valence when presented at relatively high concentrations.

**Figure 5:**
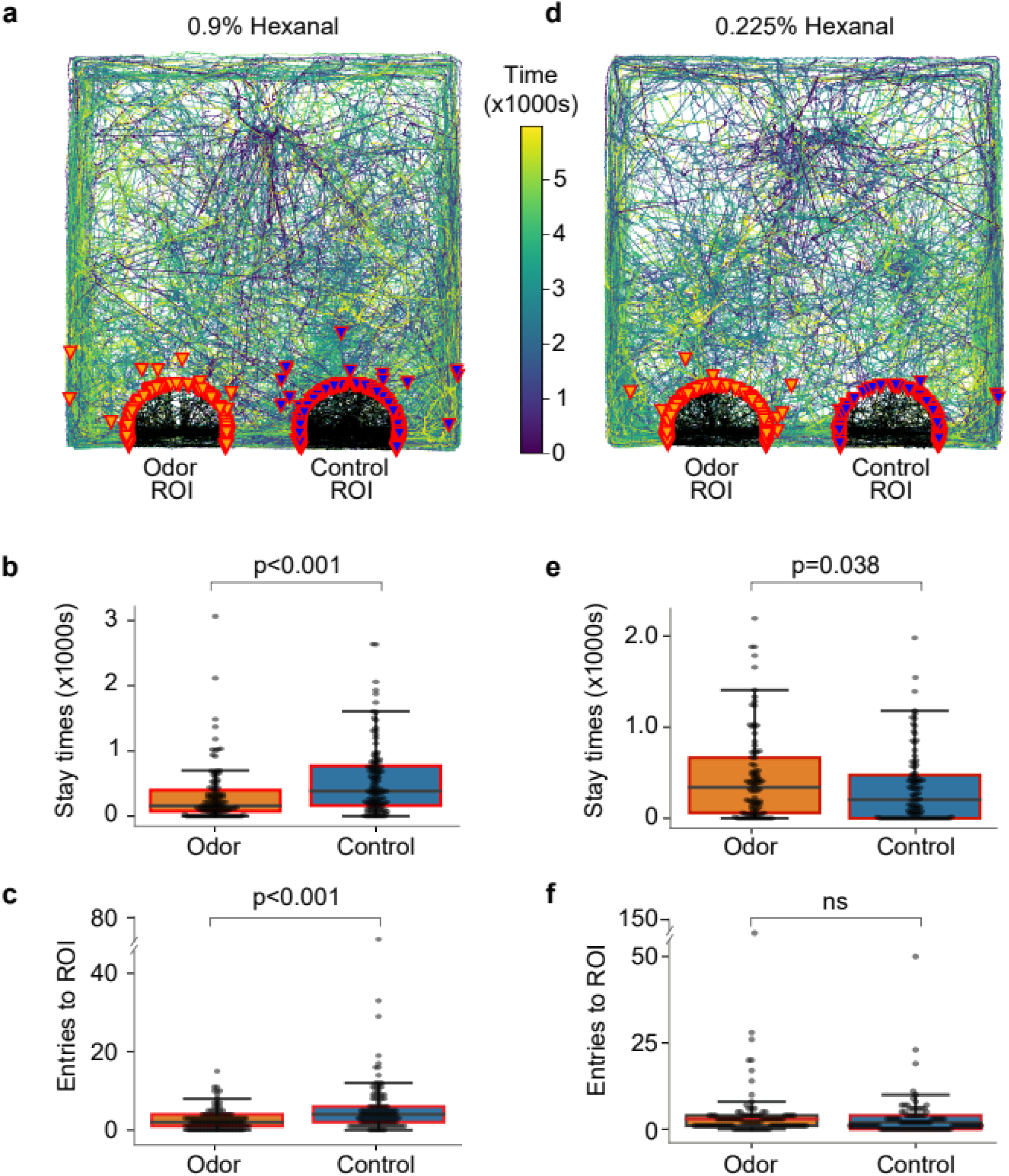
Naive hatchlings avoid a high concentration of hexanal but are attracted to a lower concentration. (a) Overlaid tracks for 0.9% hexanal, (b) comparison of stay times in the ROIs, and (c) number of entries into the ROIs. (d) Tracks for 0.225% hexanal, and comparison of (e) stay times and (f) number of ROI-entries.

### Control for mineral oil

Finally, to test whether possible odors from the mineral oil used as control in these experiments may have affected the movements of hatchlings, we repeated the 2-choice test with air passing through a clean, empty bottle tested against air passing through a bottle containing mineral oil. Hatchlings showed no preference for either choice in these experiments in terms of stay time (n = 81, with two-sided paired t-test p = 0.31; PI = 0.10), or the number of ROI entries (n = 81, with two-sided paired t-test p = 0.29; PI = 0.09) (Figure 6a-c), indicating the hatchlings were neither attracted nor repelled by mineral oil.

**Figure 6:**
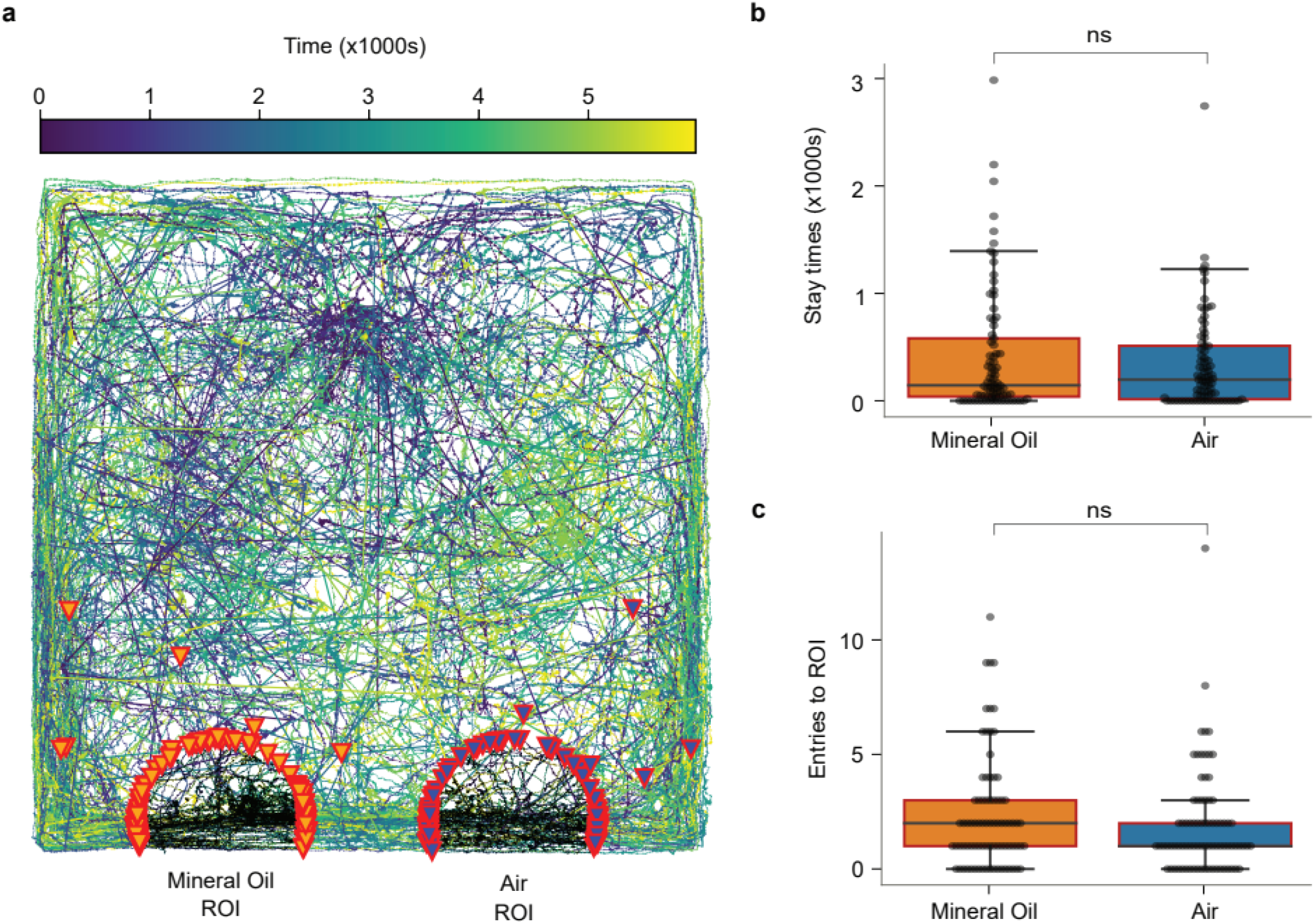
Control experiment comparing attraction to mineral oil to desiccated, filtered air. (a) Overlaid tracks of all the hatchlings, (b) comparison of stay time in the ROIs, and (c) comparison of the number of ROI entries. None of these measures showed any significant differences between filtered, desiccated air and mineral oil.

## Discussion

Here we found naïve, newly hatched locusts show an innate attraction toward the odor of wheat grass, a plant readily consumed by locusts of all ages. We also found that hatchlings require their antennae to navigate towards the odor source. What mechanism could explain this innate attraction? One possibility is hatchlings overexpress olfactory receptors sensitive to plant volatiles, allowing them to find plants by moving toward anything emitting an odor that elicits a strong antennal signal. This seems unlikely because animals are generally repelled by strong odors (Laing et al., 1978; Semmelhack and Wang, 2009; also, see figures 4-5); we are testing this idea now (Sun et al, in preparation). Another possibility is that genetically regulated hard-wiring links antennal lobe projection neurons (PNs) or glomeruli responding to food odors with neuronal pathways that assign positive valence. This, too, seems unlikely because many locust PNs distributed around the antennal lobe respond to both food odors and non-food odors (Stopfer et al, 2003; Sun et al, in preparation). Another possibility is that olfactory wiring downstream from the antennal lobe is shaped during development by volatile components of the food consumed by the mother and then deposited in egg pods. It is sometimes the case that chemical components of food, or metabolic products derived from them, function as pheromones (Ignell et al., 2001), potentially priming development. The diet of mother mice has been shown to influence olfactory neurodevelopment and odor preference in the newborn (Todrank et al., 2011). This possibility could be tested in locusts by manipulating the mother’s diet or the volatiles present in egg pods.

Our results show that even relatively low concentrations of 1-hexanol, a major monomolecular component of wheat grass that is known to elicit widespread neuronal responses in the locust olfactory pathway, evokes avoidance behavior in naïve locust hatchlings, while lower concentrations elicited neither attraction nor avoidance. This result is consistent with earlier findings in other model systems showing the importance of background odors or odor blends in determining valence. The black bean aphid, for example, is repelled by many of the individual volatile compounds of its host plant, though attracted to their blend (Webster et al., 2010). The Asian tiger mosquito is attracted to human body odor, but not to its individual components (Xie et al., 2019). And, the moth *Manduca sexta* is not attracted by single or small subsets of components present in its favored food source, the *Datura wrightii* flower, but it is attracted by larger synthetic mixtures of its major components (Riffell et al., 2009). Moreover, the odor of the *Datura* flower combined with that of the leaf of this plant elicits a stronger positive behavioral response than the floral odor alone (Kárpáti et al., 2013).

In mice, odors that separately evoke the same innate valence can evoke the opposite valence when blended (Qiu et al., 2021), a result consistent with combinatorial rather than labeled-line processing of valence information. The same logic appears to apply to our study, in which an innately attractive blend included components that evoked repulsion, suggesting that odors with innate valence are not necessarily processed by a labeled lines in locusts. Our results are consistent with earlier work in humans and in *Drosophila* showing that odor valence depends strongly on concentration (Laing et al., 1978; Semmelhack and Wang, 2009): as the concentration of an odorant increases, increasing numbers and types of odorant receptors are activated, eliciting different valence behaviors. Similarly, the convergence of excitatory and inhibitory projections from different pheromone-sensitive glomeruli in the *Drosophila* lateral horn has given rise to the suggestion that the balance of attractive and aversive signals may determine sex behaviors (Jefferis et al., 2007). A similar logic may apply for components of food odor in locusts. Further experiments testing mixtures of food odor components at different concentrations and mapping the functional pathways for these odors could help elucidate the neural basis of innate olfactory behaviors.

## Data Availability

The data underlying this article will be shared on reasonable request to the corresponding author.

## Funding

This research was supported by an Intramural Grant from NIH-NICHD to M.S.

## Acknowledgments

We thank NIMH Section on Instrumentation for help with the experimental setup, and members of the Stopfer lab, especially Zane Aldworth for helpful comments and suggestions. We also thank Erica Varga for assisting with reviewing some of the data.

## Conflict of Interest

The authors declare no conflict of interest

## Notes

### Competing Interest Statement

The authors have declared no competing interest.

